# Computations, optimization and tuning of deep feedforward neural networks

**DOI:** 10.1101/2019.12.27.889311

**Authors:** Md. Shoaibur Rahman

## Abstract

This article presents an overview of the generalized formulations of the computations, optimization, and tuning of a deep feedforward neural network. A small network has been used to systematically explain the computing steps, which were then used to establish the generalized forms of the computations in forward and backward propagations for larger networks. Additionally, some of the commonly used cost functions, activation functions, optimization algorithms, and hyper-parameters tuning approaches have been discussed.

## 1. Introduction

Deep learning is a process of training deep neural networks (DNNs) [LeCun et al. 2015]. One of the classes of DNNs is a feedforward network, which is represented with layers of computing units or activation units [Svozil et al. 1997]. Specifically, feedforward networks comprise three types of layers: an input layer, generally several hidden layers, and an output layer (**Fig. 1**). The input layer feeds the input features/data to the first hidden layer, which feeds the next layer and the process continues until the last hidden layer feeds the output layer. The input layer or layer 0 is usually not considered as an actual layer. So, with that definition, an L-layer neural network consists of L-1 hidden layers and one output layer (L^th^ layer). DNNs usually have more than one hidden layer, so they are named as the “deep” networks (depth increased by adding more hidden layers). Each hidden layer consists of several activation units or hidden units (represented by circles in figure 1), and their outputs are termed as activation values. The activation value of a unit in a layer is the input to one or more units in the next layer. In contrast, the output layer consists of one (for binary class labels) or multiple activation units (for multiple class labels). The converted class of the activation value of the output layer is an estimate of the actual class label.

**Figure 1.**
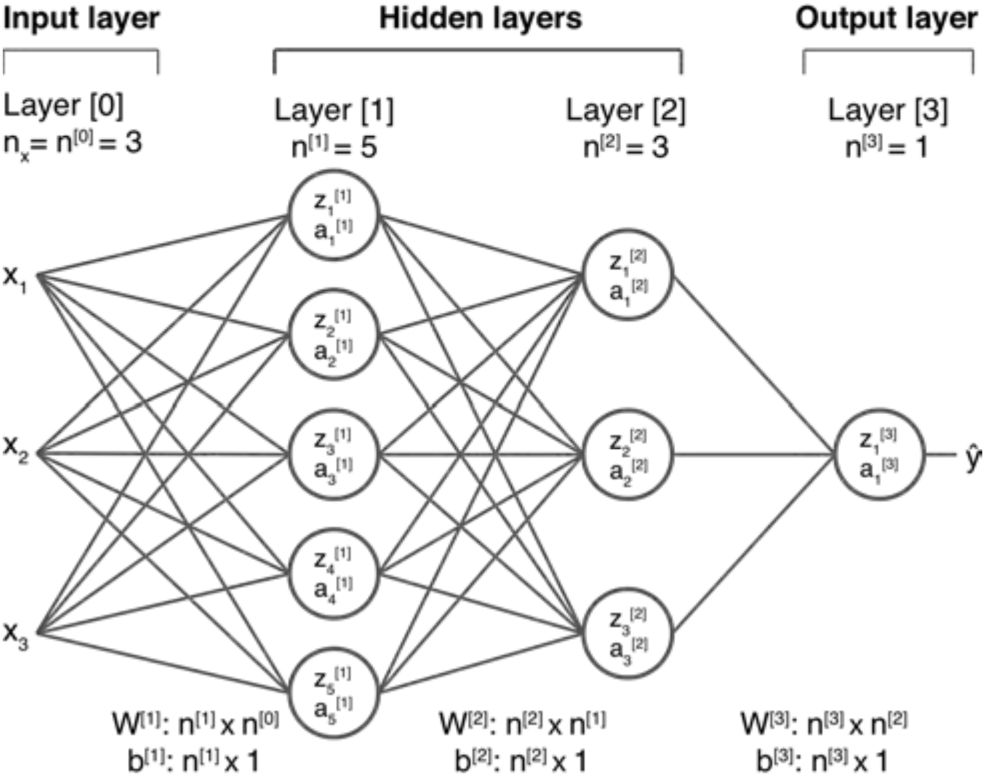
Example of a 3-layer neural network. Layer 0, the input layer is not counted as a layer. x is the input data with a number of input features n_x_ = n^[0]^ = 3. Layer 1 is the first hidden layer with *n*^[1]^ = 5 activation units (circles). Layer 2 is the second hidden layer with *n*^[2]^ = 3 activation units. Layer 3 is the output layer with *n*^[3]^ = 1 activation unit. Activation units have their respective linear transformation activation values (z and a, respectively). Activation value ŷ = a^[3]^ at the output layer is an estimate of the actual output label y. The shape of the parameter (W) at a layer is determined by the number of activation units in that layer and in the preceding layer. The number of activation units in the layer determines the shape of parameter b.

The computation in each activation unit occurs in two steps. First, the inputs to a unit are linearly transformed (like as in linear regression). For instance, if the inputs to a unit are: a_1,_ a_2,_ … a_n_, and their respective weights are: w_1_, w_2,_ … w_n_ then the transformed value z can be computed as:

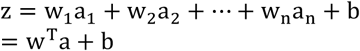

where, a and w are the vectors of the inputs and the weights, respectively, and b is a scalar that represents bias value for the activation unit. Second, the value of z is used to compute the activation value a. An activation function g is required to compute the activation value, i.e., a = g(z). Some of the commonly used activation functions are described in section 2.5.

This 2-step computation process of each unit from the inputs to the hidden layers to the output layer is termed as forward propagation. The final output of the forward propagation (i.e., activation value of the output layer), is compared with the actual output to compute an error of the estimation. This error is fed back through the hidden layers so the weights and biases can be modified or updated accordingly to adjust/minimize the error. Later in this article, we will see that the weights and biases are updated based on the derivatives of an error/cost function. The process of this feeding back of the error derivatives is termed as backward propagation or backpropagation [Johansson et al. 1991; Yu et al. 2002]. With this brief explanation, one could envision that a lot of computations are involved in both forward and backward propagations. The amount of the computations explodes if the number of the layers and the activation units increase substantially. So, understanding these computations is the essence for understanding a DNN.

In this article, we used a small 3-layer neural network to establish a general method of computations in a deep feedforward neural network. In section 2, we explained computations for forward propagation step-by-step from the input layer to the output layer and backpropagation from the output layer to the input layer. This systematic description was used to establish a general method to compute forward propagation, backpropagation, and derivatives at any given layer of a larger network. We also presented some of the commonly used cost functions and activation functions. In section 3, we discussed some techniques for the optimization of the parameters, including initialization and common algorithms. In section 4, we briefly described some of the approaches used for hyperparameter tuning. Some potential applications of DNN had been discussed in section 5. Finally, the article had been summarized in section 6.

## 2. Computations

In this section, we discuss the computations that take place in forward and backward propagations, parallelization or vectorization of the computations, and some commonly used cost functions and activation functions.

### 2.1. Forward propagation

Forward propagation starts from layer 0 (input layer) and is initiated by the input features x = a^[0]^, where the superscript number in the brace indicates the layer. Each unit of layer 1 performs a 2-step computation; first, input features are transformed linearly to compute transformation values z, and second, the transformation values are used to compute the activation values a using a given activation function g. This two-step computation is performed in layer 1 and can be represented by equations (1a-b):

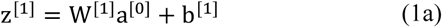

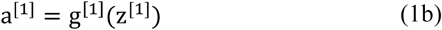

The linear transformation and activation values computed by each unit in a layer are fed into the next layer for similar computations (equations 2a-b). This forward propagation of the activation values undergoes up to the output layer (equations 3a-b). Using similar logic of linear transformation explained in the introduction, the forward propagation can be summarized as:

In layer 2:

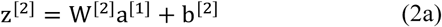

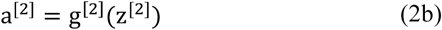

In layer 3:

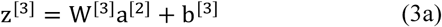

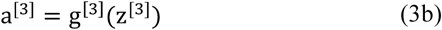

So, in an L-layer network, the general form of forward propagation at any layer l = 1, 2, … L can be expressed as:

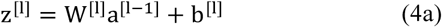

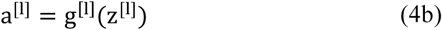

where, W is a matrix of shape (n^[1]^ × n^[l−1]^) and b is a matrix of shape (n^[1]^ × 1). The activation value a^[3]^ at the output layer is an estimation of the actual output. A cost function *𝒞* is defined based on the estimated and actual outputs (see section 2.4 for details). The costs propagate backward through the network to adjust the parameters accordingly. This loop of forward and backward propagation and parameter updates continues until the activations of the output layer get as close as possible to the actual outputs. The process of backward propagation and parameter updates will be discussed in the subsequent sections of this article.

### 2.2. Backpropagation

The goal of the backpropagation is to update the parameters at each layer based on the cost function, specifically the derivatives of the cost function with respect to the weight matrices and biases at each layer. The computations of backpropagation derivatives start from the output layer followed by at its preceding layers sequentially and stop at the first hidden layer. The calculus of partial derivatives and chain rule will be used to compute backpropagation derivatives in this article (see [Shoaib et al. 2010; Rahman 2011; Rahman and Haque 2012] for some advanced applications of partial derivatives and chain rule). For simplicity, the partial derivative of the cost function with respect to any variable ϕ will be denoted as Δϕ, i.e.,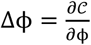.

Derivatives of the cost function with respect to the weight matrix and bias vector in layer 3 can be computed using the chain rule, i.e.,

In layer 3:

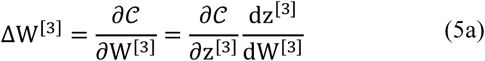

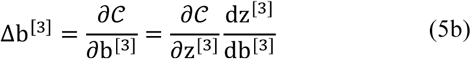

Now, using chain rules:

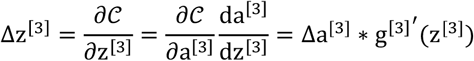

where, 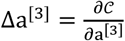 and g^[3]^′(z^[3]^) are the derivatives of the activation function in layer 3 (see section 2.5 for the details of the activation functions) and * indicates an element-wise product.

Also, from equation (3a), it can be shown that:

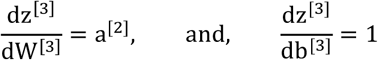

Therefore, the evaluation of the derivatives of the cost function with respect to the parameters in layer 3 can be summarized using equations (6a-d):

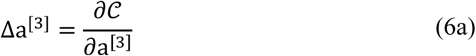

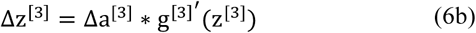

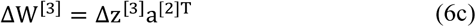

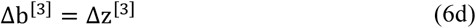

Similar approach can be applied in layer 2:

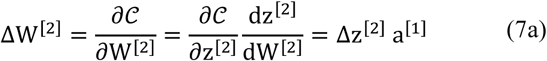

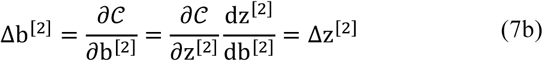

where,

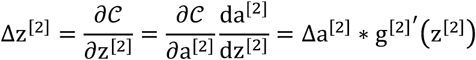

and,

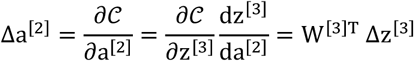

Therefore, the evaluation of the derivatives of the cost function with respect to the parameters in layer 2 can be summarized using equations (8a-d):

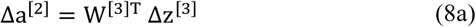

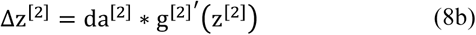

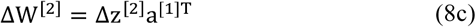

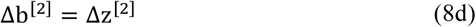

Finally, similar approach can be applied in layer 1:

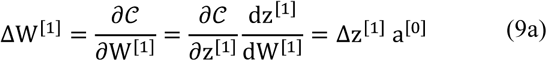

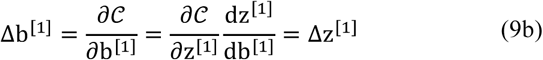

where,

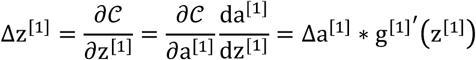

and,

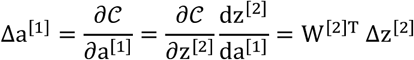

Therefore, the evaluation of the derivatives of the cost function with respect to the parameters in layer 1 can be summarized using equations (10a-d):

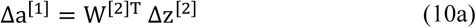

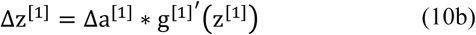

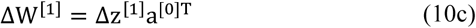

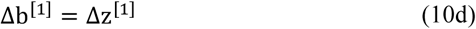

Since, Δb is a single value at a given layer, all the elements of Δz are summed to estimate Δb during the implementation, i.e., Δb = ∑Δz.

Using equations (6, 8, and 10), the computations of backpropagation derivatives at any layer can be generalized as:

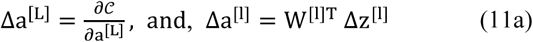

where, l = L − 1, L − 2 … 1

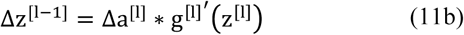

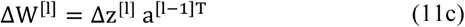

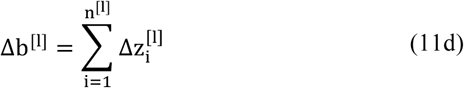

So far, only one input variable, x = a^[0]^ of features n_x_ = n^[0]^, has been used in the forward propagation and backpropagation. So, the procedures can be looped m times with m features or training examples, but this may substantially increase the computation time as m gets larger (usually m is very large in deep learning problems). An alternative of looping is to use a matrix computation approach, which will vectorize (or parallelize) the computations for all training examples together, and thus will make the computations faster. The next section focuses on the vectorized implementation of the forward and backpropagation.

### 2.3. Vectorization of propagation

To vectorize the computations, an input matrix X is formed using m training examples with each example forming one column of X. Input matrix X will also be denoted as A^[0]^ to generalize the equations for any given layer. So,

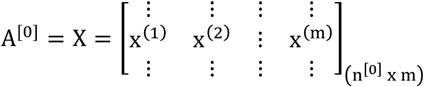

where, 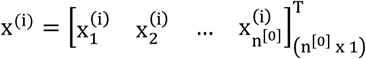, i = 1 … m

Similarly, for a given layer l, matrices Z and A can be formed as follows:

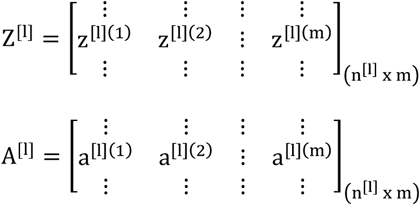

where, 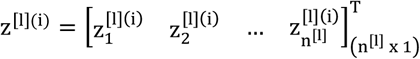 and 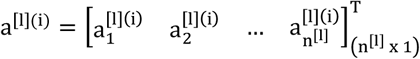, i = 1 … m

Additionally, each training example will have one output label, so the output matrix Y will have a shape of (1 × m), i.e.,

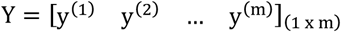

However, the shape of the parameters W and b remain unchanged, as they are independent of the number of training examples. Given these matrices setup, equations (4a-b) can be turned into matrix forms to vectorize the forward propagation, i.e.,

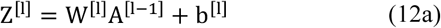

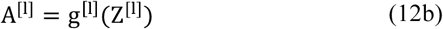

The outputs from the forward propagation are used to compute the cost. In vectorized computations, each training example will contribute to the cost. So, the total cost will be the summation of costs contributed by all training examples. This total cost is used to initiate the backpropagation from the output layer.

Using the same matrix setups, equations (13a-d) can be turned into matrix forms to vectorize the backpropagation, i.e.,

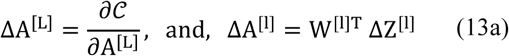

where, l = L − 1, L − 2 … 1

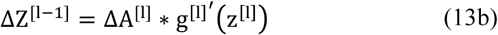

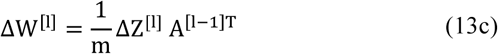

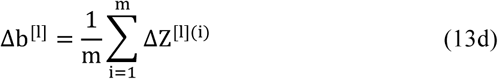

The backpropagation starts from the output layer and is initiated by ΔA^[L]^, which depends only on the cost function, unlike in hidden layers. The derivatives of the parameters are scaled by m, which implies that an average contribution from all training examples is used to update the parameters.

This is obvious from equations (12-13) that the cost functions and activation functions are required to optimize the parameters of a particular network when an input/output dataset is given, the cost function and activation functions are known. Importantly, activation functions are required for forward propagation (equation 12b) and its derivatives are required for backpropagation (equation 13b). Moreover, the same activation function can be used for all layers, or different activation functions can be used for different layers. The choice for the activation functions at different layers is also a part of hyperparameter tuning. The following subsections briefly describe some of the commonly used cost and activation functions in deep learning.

### 2.4. Cost functions

Cost function, or more specifically its derivative, is required to initiate the backpropagation at the output layer (equation 13a). Some commonly used cost functions in deep learning are mean squared error, mean absolute error, hinge loss, squared hinge loss, categorical cross-entropy loss (for multi-class classification) and binary cross-entropy loss (for binary classification). Here, we will explain only binary cross-entropy cost function, which is a widely used cost function for binary classifications using deep learning.

Given i^th^ training example x^(i)^, the probability of binary output y^(i)^ is defined as:

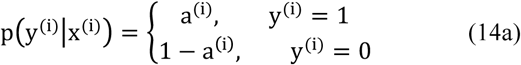

Alternatively,

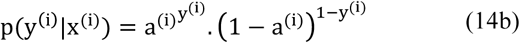

Thus, for m independent variables, the likelihood or log-likelihood are defined, respectively, as:

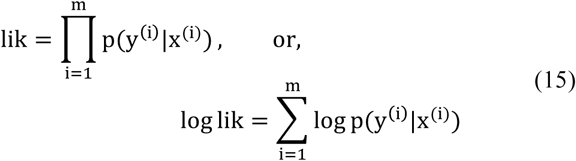

To maximize log-likelihood, we minimize the negative loss function

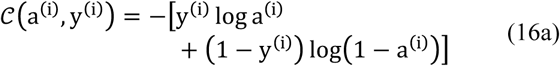

As explained in the previous sections, the cost function is formed based on the actual output Y and the estimated outputs A^[L]^ at the output layer. So, equation (16) can be turned into a matrix form to establish the cost function as:

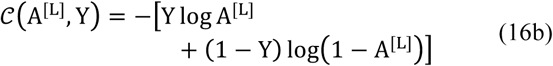

The derivative of the cost function with respect to A^[L]^ using simple differential rule of calculus can be computed as:

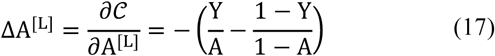

This derivative will be used in equation (13a) to initiate the backpropagation.

### 2.5. Activation functions

Unlike cost function, activation functions are required for forward propagation (equation 12b) and its derivative for backpropagation (equation 13b). Some of the commonly used activation functions in deep learning are sigmoid, softmax, hyperbolic tangent (tanh), rectifier linear unit (ReLU) and leaky ReLU. Their general definitions, derivatives and usages are briefly explained in this section. Also, a summary is given in Table 1 and graphical visualizations are represented in Figure 2.

**Table 1.**
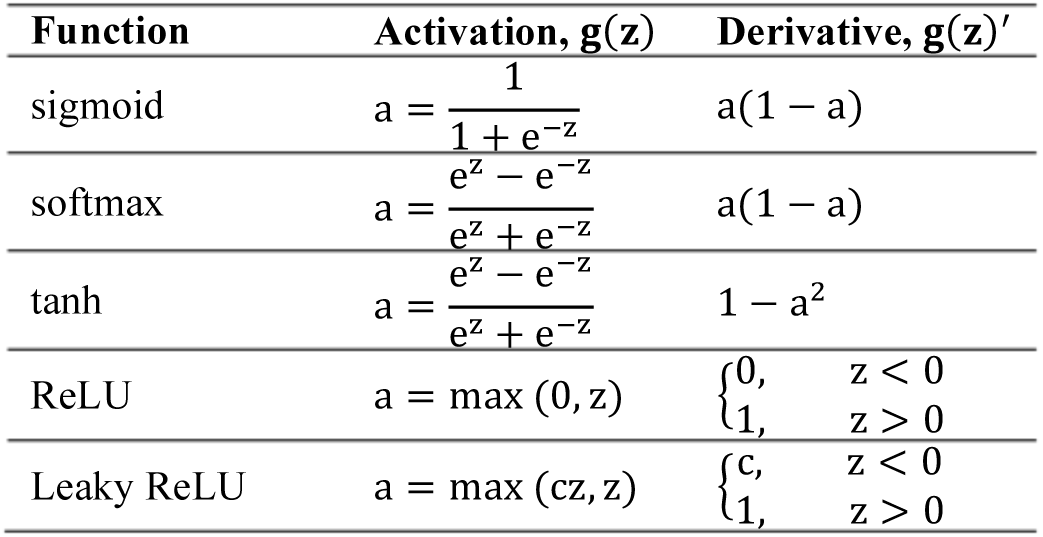
Activation functions and their derivatives

| Function | Activation, $g(z)$ | Derivative, $g(z)'$ |
| --- | --- | --- |
| sigmoid | $a = \frac{1}{1 + e^{-z}}$ | $a(1 - a)$ |
| softmax | $a = \frac{e^z - e^{-z}}{e^z + e^{-z}}$ | $a(1 - a)$ |
| tanh | $a = \frac{e^z - e^{-z}}{e^z + e^{-z}}$ | $1 - a^2$ |
| ReLU | $a = \max(0, z)$ | $\begin{cases} 0, & z < 0 \\ 1, & z > 0 \end{cases}$ |
| Leaky ReLU | $a = \max(cz, z)$ | $\begin{cases} c, & z < 0 \\ 1, & z > 0 \end{cases}$ |

**Figure 2.**
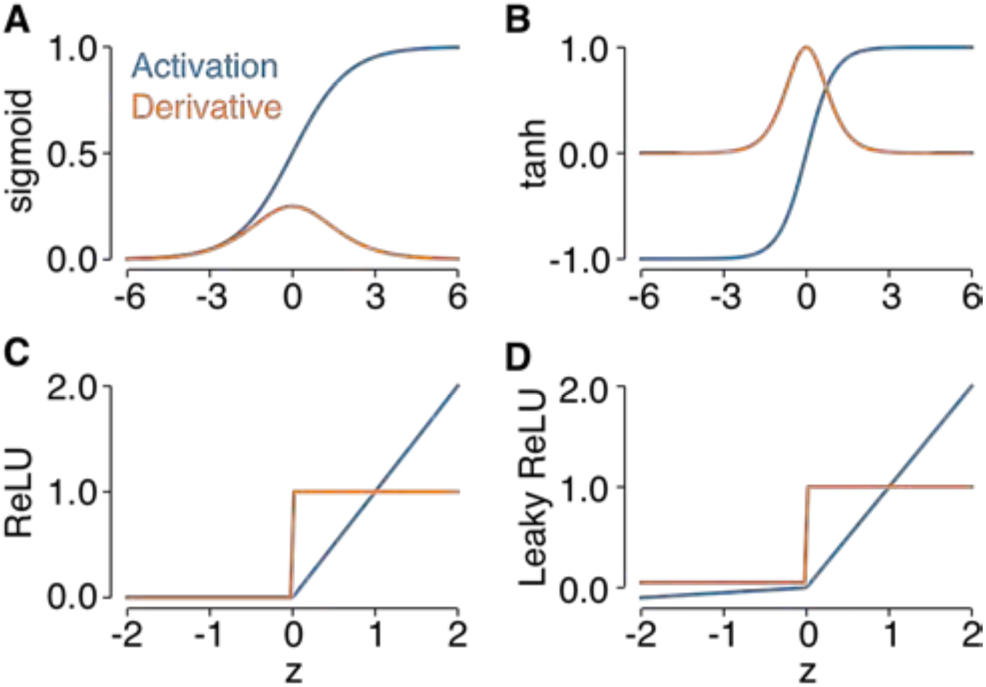
Activation functions and their derivatives. **A**) sigmoid. **B**) Hyperbolic tangent. **C**) ReLU. **D**) Leaky ReLU with a constant value of 0.05. Y-axis represents both activations and derivatives; x-axis represents the values which activation and derivatives were computed.

Sigmoid activation:

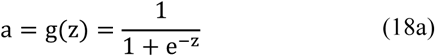

Derivative of sigmoid:

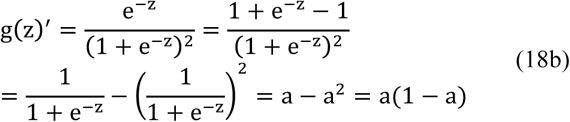

Softmax activation:

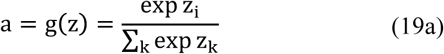

Derivative of softmax:

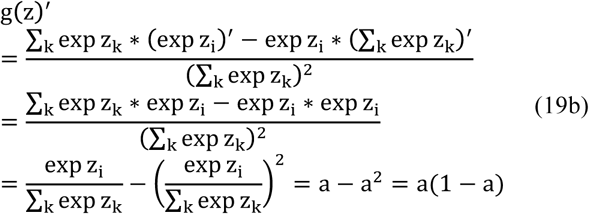

Tanh activation:

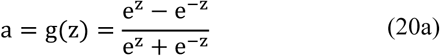

Derivative of tanh:

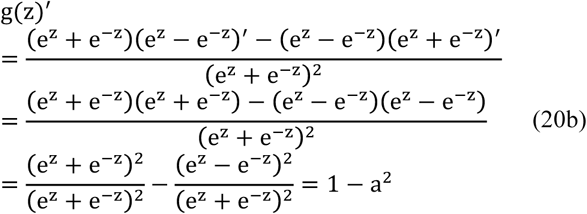

ReLU activation:

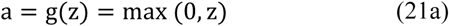

Derivative of ReLU:

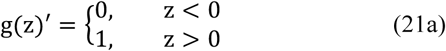

In case of leaky ReLU, a = max (cz, z), so its derivative will be a constant c for z < 0 and 1 for z > 1.

Sigmoid and softmax are used at the output layer respectively for binary and multi-class classification problems. However, neither of them is a good choice for the hidden layers. In contrast, tanh works better than sigmoid/softmax in hidden layers because the average of the outputs of tanh is close to zero, which centers the data well for the next layer. Nevertheless, for a larger absolute value of z, the derivatives of tanh becomes close to zero. Therefore, the optimization (e.g., gradient descent) slows down the learning or training of the model. To avoid these problems, ReLU is a good choice as its gradient is always 1 for positive values. Yet, for negative values, ReLU’s gradient is 0. So, leaky ReLU can be used as an alternative to ReLU. It has been shown that both ReLU and leaky ReLU perform well in practice.

## 3. Parameter optimization

Deep learning models comprise a lot of parameters, sometimes exceeding few millions for a moderately large model. So, optimization of the parameters is crucial for a good performance of the model. Indeed, the development of efficient methods for searching optimal parameters has been one of the major reasons for the recent success of the deep learning. These methods include parameter initialization, update rules, and optimization algorithms.

### 3.1. Initialization

Proper initialization of the parameters is crucial to achieve desired performances of the deep neural network models.

#### Zero initialization

Initialize parameters to all zeros. Although zero initialization works for small models like logistic regression, it does not work well for deep neural networks.

#### Random initialization

Initialize parameters to all random values drawn from a distribution, e.g., uniform or normal distributions. It works better than zero initialization. However, random initialization potentially leads to two issues: vanishing gradients and exploding gradients.

When the weights are initialized to very small values, the gradients of costs tend to zero. So, the parameters get updated very slowly, or in worst case, it stops updating the parameters. This phenomenon is known as vanishing gradient. Conversely, when the weights are initialized to larger values, they get multiplied along the layers so cause a larger change in the cost. So, the gradients also get larger. This phenomenon is known as exploding gradients.

To avoid problems of vanishing and exploding gradients, the randomly generated values can be multiplied by a small factor. The multiplication factor could be a small constant (e.g., 0.01), or can be determined based on the number hidden units for a given layer, as suggested below:

#### Xavier initialization [Glorot and Bengio 2010]

For a given layer l, the randomly generated values are multiplied by 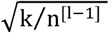, where k = 1 or 2, for ReLU or tanh activation functions, respectively, and n^[l−1]^ is the number of hidden units in previous layer, i.e., in layer l − 1. Xavier initialization is also termed at Glorot initialization.

#### He initialization [He et al. 2105]

For a given layer l, the randomly generated values are multiplied by 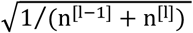, where n^[1−1]^ and n^[l]^ is the number of hidden units in layers l and l − 1.

### 3.3. Optimization algorithms

The eventual goal of the forward propagation and backpropagation is to optimize the parameters at each layer of the network. In other words, the parameters are updated based on the backpropagation derivatives during every cycle (iteration) of the propagation. The update rules depend on which optimization algorithm is in use [Ruder 2016]. This article focuses on some of widely used algorithms, e.g., gradient descent (mini-batch and stochastic gradient), adam, momentum, and RMSProp optimizer.

#### Gradient descent optimizer

The gradients of the cost function with respect to the parameters are computed, which are then used to update the parameters as follows:

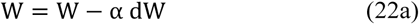

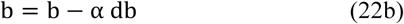

where, W and b are the weight and bias parameters, dW and db are the derivatives of the cost function with respect to the weights and biases, respectively, and α is called the learning rate, which can be tuned to improve the performance of the model. Sometimes gradient descent is termed as batch gradient descent, specifically when this is applied on the full training set simultaneously. Two other variations of this optimizer are mini-batch and stochastic gradients:

#### Mini-batch gradient descent

In many applications of deep learning, the number of training examples is large. In such cases, the training examples are divided into mini-batches (small groups), and gradient descent is applied on each mini-batch sequentially, which is known as mini-batch gradient descent optimization.

#### Stochastic gradient descent

Sometimes the model is trained using each training example, and thus gradient descent is applied to each training example sequentially. This optimization procedure is known as stochastic gradient descent. This is useful for online learning.

#### Adam optimizer [Kingma and Ba 2015]

Adam optimization method computes individual adaptive learning rates for different parameters from estimates of first and second moments of the gradients. The contributions from the first and second moments are estimated using exponentially moving average approach. The first moments (i.e., the means) v_dw_ and v_db_ are estimated as:

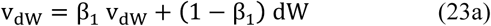

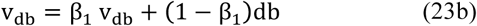

with bias-corrected moments as:

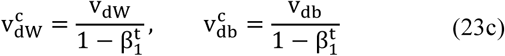

And second moments (i.e., the uncentered variances) s_dw_ and s_db_ are estimated as:

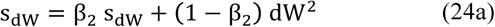

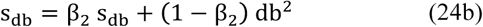

with bias-corrected moments as:

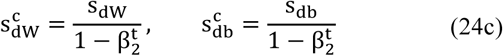

Using these corrected moments, the parameters are updated using the following rules:

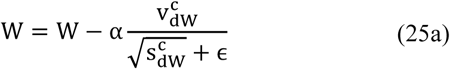

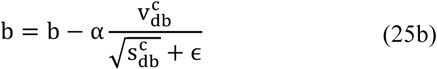

where, β_1,_ β_2,_ t, and ϵ are the hyper-parameters and need to be tuned.

#### Momentum [Qian 1999]

When only the first moments are considered, then the optimization is known as momentum (or gradient descent with momentum). The parameters are updated using the following rules:

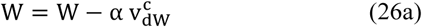

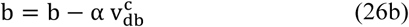

#### Root Mean Squared Propagation (RMSProp) [Tielman and Hinton, 2012]

When only the second moments are considered, then the optimization is known as RMSProp. The parameters are updated using the following rules:

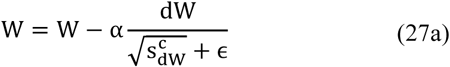

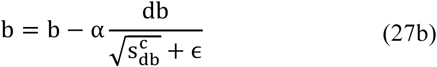

The details of these algorithms as well as other types of optimization techniques can be found in [Ruder 2016].

## 4. Hyper-parameter tuning

In this section, we discuss two things: a set of the most common hyper-parameters used in feedforward networks, and the approaches to tune those hyper-parameters.

### 4.1. Common hyper-parameters

The most common hyper-parameters used in feedforward networks are related to network architectures, regularization, optimization and training.

The most common hyper-parameters related to the network architectures include the number of hidden layers and the number of hidden units in each layer. Sometimes, different activation functions can be used at different layers, although as discussed previously, ReLU or leaky ReLU in the hidden layers works well almost always.

The most common hyper-parameters related to regularization include L2 regularization [Tikhonov,1943], lasso [Tibshirani 1996], dropout [Srivastava et al. 2014] and batch normalization [Ioffe and Szegedy 2015]. Regularization prohibits overfitting of the model to the training set.

According to the L2 regularization, the regular cost is penalized by the L2-norms of the parameters, so the total cost increases to:

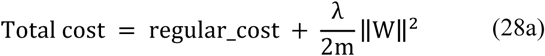

Therefore, the derivative of the total cost with respect to the weight parameter becomes:

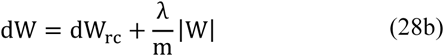

where, dW and dW_rc_ are the derivatives of the total cost and regular cost, respectively, m is the number of training examples, and λ ≥ 0 is a regularization hyper-parameter. Common practice is to train the model with logarithmically spaced values of λ.

Conversely, according to the lasso or L1 regularization, the regular cost is penalized by the L1-norms of the parameters, so the total cost increases to:

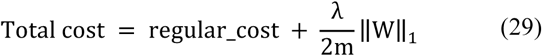

Note that the total cost in this case has no derivative at the origin because L1-norm is not differentiable at the origin. Therefore, the traditional derivative-based optimization method like gradient descent cannot be used. Instead, coordinate descent method can be useful to optimize this total cost.

Dropout is a simple technique to prevent neural networks from overfitting. It is simple in the sense that some connections between the hidden units are randomly dropped [Srivastava et al. 2014]. Therefore, the complexity of the network reduces and the network becomes more linear as the dropout increases. The amount of dropout, i.e., how many connections or what fraction of the total connections to be dropped, can be tuned to achieve the best performing model.

According to the batch normalization [Ioffe and Szegedy, 2015], for each batch (more commonly known as mini-batch), the transformation values are normalized before they are used to compute activation at each layer, i.e.,

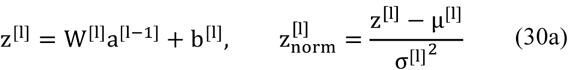

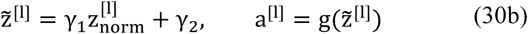

where, µ^[I]^ and σ^[l]^ are respectively the mean and standard deviation of z^[l]^. Using hyper-parameters γ_l_ and γ_2_, we can transform 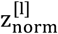 to any value 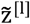 with a variance (γ_l_) and a mean (γ_2_) of what we want.

Two things to note about batch normalization: first, since z^[l]^ is normalized, the constants b^[l]^ do not contribute to the computations of a^[l]^. So, parameter b^[l]^ can be ignored for batch normalization. Second, the test set should be normalized using µ^[I]^ and σ^[l]^ values obtained from the training sets.

The most common hyper-parameters related to the optimizers are the learning rate and learning rate decay. Learning rate determines how much the parameters will change after each iteration. Common practice is to train the model with logarithmically spaced values of learning rate. Additionally, learning rate can be changed after each or some iterations by using learning rate decay hyper-parameter. The types of other hyper-parameters depend on what optimization algorithm is used. For adam optimizer, the common hyper-parameters are first moment parameter (β_1_), second moment parameter (β_2_), bias correction parameter (t) and numerical stability parameter (ϵ). The default values in different platforms are: β_1_ = 0.9, β_2_ = 0.999 and ϵ = 1e^−8^, but tuning this hyper-parameters can improve the performance in many applications.

The most common hyper-parameters related to the training of the model include mini-batch size and epoch number. The mini-batch size is usually chosen so it becomes a power of 2, e.g., 16, 32, 64 etc.

### 4.2. Tuning approaches

Several approaches have been suggested for hyper-parameter tuning [Begstra et al. 2001], but there is no absolute guideline about exactly which approach to use. Commonly used approaches are grid search, random search [Bergstra and Bengio 2012], and Bayesian optimization [Snoek et al. 2012]. Here we discuss the grid search and the random search approaches.

#### Grid search

A grid is formed using the subset of values of all hyper-parameters. The model is trained using each set of values for each grid point. The set that provides the lowest training cost is used to test the model with the test data. However, some hyper-parameters may not be important for performance improvement. In grid search, we train the model with many of these less important values, which makes the searching less efficient. So, grid search is not a good choice unless we have a small number of hyper-parameters.

#### Random search

The problem of grid search can be avoided by using random search, where a set of values of all hyper-parameters are randomly chosen, on which the model is trained. This process is performed for several times, and the set of values that provides the lowest training cost is used to test the model with the test data. Random search usually performs better than a grid search.

## 5. Applications

DNNs are commonly used to classify data in different fields. For example, convolutional neural networks (CNNs) in addition to feedforward networks have widely been used in computer vision applications, e.g., image classification and recognition [LeCun et al. 1998; Krizhevsky et al. 2012; He et al. 2016]. Recurrent neural networks (RNNs) have been used in natural language processing [Hirschberg and Manning 2015; Socher et al. 2011; Young et al. 2018], e.g., text classification [Lai et al. 2015] and speech recognition [Graves et al. 2013]. Moreover, recent studies show that DNN has tremendous potential in neuroscience [Marblestone et al. 2016; Güçlu and van Gerven 2014] and psychological applications [Choi et al. 2018]. Yet, it remains to uncover how DNNs relate with brain functions and psychology. However, further exploration of variety of psychophysical studies [Rahman et al. 2017; Convento et al. 2018; Rahman and Yau 2019; Rahman 2019] and functional neuroimaging experiments [Rahman et al. 2019; Kay et al. 2008; Pereira et al. 2018; Huth et al. 2016; Kamitani et al. 2005] with DNN may bridge the gap between neuroscience and deep learning.

## 6. Discussion

This article provides the essentials of the computations in a DNN. We explained what computations take place in the hidden layers/units during forward and backward propagation. Using a small network, we systematically developed a generalized form of the computations at each layer of a network with any number of hidden layers and hidden activation units. We also presented how these computations can be vectorized to make the computations more efficient. Moreover, we briefly described some of the commonly used cost functions, activation functions, and optimization algorithms in DNN. Finally, we talked about the some the studies that brought deep learning with a tremendous success in computer vision and language processing. We also presented some emerging studies and application of deep learning in other fields, including neuroscience and psychology, which may uncover many of the important but unsolved problems in the respective fields.

The computations, parameter optimization and tuning of the hyperparameters approaches presented in this article were implemented with Numpy, TensorFlow and Keras. The codes are available in Cat_Images and Hand_Signs folders on: https://github.com/shoaibur/Deep_Learning The goal of these codes was not to provide a fine-tuned program for any particular applications. Instead, we focused on the development and implementation of a generalized method for DNNs, so users can use them to build their application-specific programs.

## Acknowledgement

Many of the terminologies in this article were used in a similar manner as they were used in Andrew Ng’s deep learning classes on Coursera.

